# Advances in deep reinforcement learning enable better predictions of human behavior in time-continuous tasks

**DOI:** 10.1101/2025.05.20.655119

**Authors:** Sabine Haberland, Hannes Ruge, Holger Frimmel

## Abstract

Humans have to respond to everyday tasks with goal-directed actions in complex and time-continuous environments. However, modeling human behavior in such environments has been challenging. Deep Q-networks (DQNs), an application of deep learning used in reinforcement learning (RL), enable the investigation of how humans transform high-dimensional, time-continuous visual stimuli into appropriate motor responses. While recent advances in DQNs have led to significant performance improvements, it has remained unclear whether these advancements translate into improved modeling of human behavior. Here, we recorded motor responses in human participants (N=23) while playing three distinct arcade games. We used stimulus features generated by a DQN as predictors for human data by fitting the DQN’s response probabilities to human motor responses using a linear model. We hypothesized that advancements in RL models would lead to better prediction of human motor responses. Therefore, we used features from two recently developed DQN models (Ape-X and SEED) and a third baseline DQN to compare prediction accuracy. Compared to the baseline DQN, Ape-X and SEED involved additional structures, such as dueling and double Q-learning, and a long short-term memory, which considerably improved their performances when playing arcade games. Since the experimental tasks were time-continuous, we also analyzed the effect of temporal resolution on prediction accuracy by smoothing the model and human data to varying degrees. We found that all three models predict human behavior significantly above chance level. SEED, the most complex model, outperformed the others in prediction accuracy of human behavior across all three games. These results suggest that advances in deep reinforcement learning can improve our capability to model human behavior in complex, time-continuous experimental tasks at a fine-grained temporal scale, thereby opening an interesting avenue for future research that complements the conventional experimental approach, characterized by its trial structure and use of low-dimensional stimuli.

**Author summary:** In our complex environment, we are constantly responding to outside influences. Traditional trial-based psychological experiments have limitations because they cannot account for the continuous nature of the world. In this study, we combine psychological questions with advanced technology to investigate how humans behave over time. Artificial neural networks have not only proven they can outperform humans in complex tasks but also have the ability to generate features that can be used to model human behavior. We investigated whether technological advancements in deep neural networks would lead to higher accuracy in predicting human motor responses recorded from subjects while playing three arcade games. We compared the predictive performance of features generated by three neural networks of varying complexity. Our results provide evidence that all three models can predict human behavior, with the most advanced model achieving the highest prediction accuracy. These results suggest that advanced neural networks are suitable models for studying human behavior in a time-continuous context and serve as a complement to the trial-based study designs.

## Introduction

In their daily lives, humans are confronted with tasks in complex and time-continuous environments, leading to the development of the ability to quickly react to high-dimensional information with goal-directed actions. While the conventional experimental approach, characterized by its discrete structure and use of low-dimensional stimuli, has offered important insights into human behavior, modeling human behavior in time-continuous environments has been challenging.

In recent years, a new approach has emerged: predicting human behavior using a generalized linear model (GLM) with features generated by a deep Q-network (DQN). DQNs have been created through the use of deep learning models in the field of reinforcement learning (RL) [1] and have been shown to be suitable models for addressing research questions [2] within cognitive neuroscience. They have enabled the investigation of how the brain processes time-continuous tasks by parameterizing the process of stimulus-response transformations. This is achieved by Q-learning [3]: In each state *s_t_*, an agent selects an action *a_t_*according to a certain stochastic policy *π* that maps states to actions, and observes a reward *r_t_*(*s_t_, a_t_*) generated by the game at time step *t* to proceed to the next state *s_t_*_+1_. For the policy *π*, the corresponding Q-function *Q^π^*(*s, a*) of the state-action pair (*s, a*) is defined as the expected, discounted future reward:

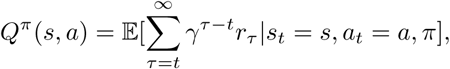

with the discount factor *γ* ∈ [0, 1], which determines the importance of future rewards. Deep Q-learning allows the agent to estimate the Q-values and to approximate the optimal policy *π*^∗^ that maximizes the expected, discounted future reward by primarily following the strategy of selecting the action with the highest Q-value. There is preliminary evidence that human gaming behavior can be modeled using features of a basic DQN [4]. Beyond modeling human behavior, a basic DQN model can also be used to predict human brain activity [5].

The ongoing rapid improvements in the field of RL have led to more advanced DQN models that solve complex tasks at human-level and beyond [6, 7]. Based on the previous results, the question arose as to whether advancements in machine learning can be leveraged to enhance the modeling of human behavior. Arcade games serve as a suitable benchmark for evaluating the performance of DQNs and for studying time-continuous stimulus-response (S-R) transformations with high-dimensional visual stimuli, as they typically demand immediate actions. Here, we consider three different DQNs: a baseline DQN [8], Ape-X [9], and SEED [7]. All three DQNs possess the capability to acquire the skill of playing arcade games through deep Q-learning. They differ in their structure, training, and model complexity. The baseline DQN is a “vanilla” feed-forward convolutional neural network (CNN) combined with a Q-learning algorithm. The CNN comprises an input layer, followed by a series of hidden layers, namely convolutional layers and fully connected layers, each with a non-linear activation function, and an output layer. Each neuron of the output layer is assigned to a possible action of a game and thus indicates the Q-value associated with the action for an input state. While Ape-X and SEED share some structure with the baseline DQN, they incorporate additional elements in architecture and learning methods that lead to a significant increase in performance. In the case of Ape-X and SEED, a dueling architecture has been integrated. The estimation of Q-values is split into two components: one stream for the evaluation of different actions and another for evaluating a certain state. Moreover, SEED introduces a substantial enhancement to its architecture by implementing a long short-term memory (LSTM), a recurrent neural network (RNN) that incorporates past experiences into decision-making. Ape-X and SEED have implemented additional techniques beyond their architecture to enhance Q-learning. For instance, they employ double Q-learning by utilizing separate networks for action evaluation and action selection. Additionally, they collect more gameplay data, evaluate the data according to its importance, and store it in a shared buffer. Unlike the other two DQNs, SEED does not suffer from the limitation of reward clipping, restricting the received reward to a certain range. We individually trained the baseline DQN and Ape-X to play arcade games and used a pre-trained SEED. The baseline DQN, compared to the other two DQNs, does not show stable learning curves. After sufficient training, Ape-X and SEED achieved performance significantly higher than human-level. In contrast, despite its basic architecture, the baseline DQN achieved game scores close to human levels but slightly underperformed them.

We analyzed behavioral data recorded from subjects playing the arcade games Breakout, Space Invaders, and Enduro. In Breakout, the player must control a paddle to smash a wall of bricks with a ball. In Enduro, a racing game, the player aims to overtake as many cars as possible across different track conditions. In Space Invaders, the player must fight aliens with a spaceship before they reach Earth. These games were selected to encompass a wide range of game types and skills.

We used the trained DQN models as a nonlinear, feature-generating mapping by processing the video screens seen by subjects through the DQNs. We used the features generated by DQNs to predict human responses using a GLM, which fitted the Q-values to human data. We hypothesize that the improvements in performance of Ape-X and SEED translate into better prediction accuracy of behavioral data in the form of human motor responses.

To test this hypothesis, we evaluated and compared the prediction accuracy of the linear model using features of Ape-X and SEED against the linear model using features of the baseline DQN. Additionally, as the experimental tasks were time-continuous, we analyzed the impact of temporal resolution on prediction accuracy by smoothing DQN and human data to varying degrees. We also explored whether an adequate amount of training time for the DQNs is essential for achieving high prediction accuracies. We successfully showed that GLMs with features from a trained DQN are suitable models for predicting human behavior at a fine-grained temporal scale.

## 1 Results

We investigated the behavior of 23 human participants (15 female, 8 male; mean age: 24.7 years, age range: 20-34 years) playing three arcade games: Breakout, Space Invaders, and Enduro. We recorded gaming data, including motor responses, gameplay videos, which were sequences of observed screens, and the corresponding rewards. The task and the collected data were sampled at 45 Hz. From a technical perspective, the implementation of the task was discrete due to the presentation of individual frames, although from a human perceptual standpoint, the gameplay appeared time-continuous [10]. The participants played each of the games for 7 sessions, with each session lasting 7 minutes of gameplay after they had practiced the game. The analysis consisted of two steps (see Fig 1).

**Fig 1.**
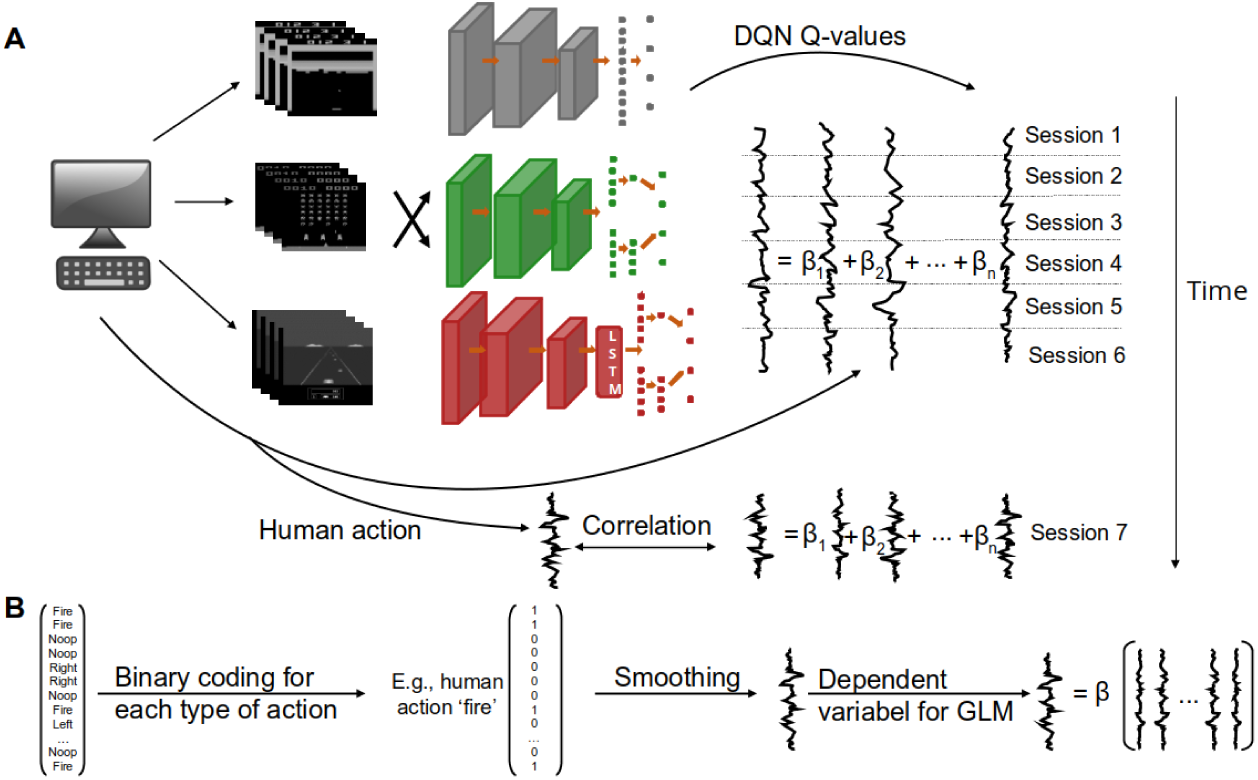
DQN-based linear model. (A) Behavioral data were collected while subjects played the arcade games Breakout (top), Space Invaders (middle), and Enduro (bottom). The human-generated videos were processed by a baseline DQN (gray), Ape-X (green), and SEED (red). Each neuron of the output layer of a DQN corresponds to the Q-value of a possible type of action, such as ‘no operation’ (’noop’), move left, move right, fire, and brake. The time series of Q-values were used as predictors in a GLM, while the actual time series of human motor responses served as the predicted variable. The GLMs were fitted to six out of seven sessions, and the Pearson correlation between the predicted human time series and the actual one was calculated on the left-out session. (B) In the preprocessing step, human actions were binary coded and smoothed. A separate GLM was fitted for each type of human action, with the corresponding human time series as the dependent variable.

In the first step, the DQN models were used for feature generation. Preprocessed human-generated videos of a particular game were processed by the DQNs trained on this game, resulting in a time series of activation values for each neuron in the top layer. Each neuron within the output layer represents a possible action in the game, with the activation serving the Q-value of a video frame. These time series were then used as predictors in a GLM to map the stimulus features to human actions. Before their application in a GLM for predicting human data, the human and DQN time series had to be preprocessed, including smoothing using a Gaussian kernel. For modeling human motor responses, a smoothing kernel with a full width at half maximum (FWHM) of 0.79 seconds proved to be optimal (see Section 1.2), which was used for smoothing in Sections 1.1 and 1.3. Further details can be found in Section 3.2.4, and a visualization of these time series is presented in S1 Fig. In the second step, for each DQN, each game, each subject, and each type of action we employed a separate logistic model to fit the generated features to human data. The time series of Q-values served as predictors, while the time series of human actions was used as the dependent variable. To evaluate the prediction accuracy, a 7-fold cross-validation procedure was employed. We calculated the Pearson correlations between the preprocessed, actual human time series and the predicted time series as mean values across all test blocks, subjects, and types of actions. In the following, when we refer to the prediction accuracy of a DQN, we are specifically referring to the prediction accuracy of the GLM model, which utilizes predictors generated by the DQN.

### 1.1 Improvements in DQNs contribute to improved predictive power

Advancements in deep RL have significantly enhanced the performance of DQNs in playing arcade games. SEED and Ape-X have achieved levels of performance surpassing those of humans [7, 9]. However, it remained unclear whether the more advanced DQNs generate features for the GLM that yield higher correlations compared to the baseline DQN. Therefore, we compared the prediction accuracy of human motor responses.

The results show that all three models predicted human behavior significantly above chance level (see Table 1) in all three games.

**Table 1.** Results of the one-sample t-test.

|  | Baseline DQN | Ape-X | SEED |
| --- | --- | --- | --- |
| Breakout | $p < .001$, $t(20)=29.19$,<br>$d=6.37$ | $p < .001$, $t(20)=28.82$,<br>$d=6.29$ | $p < .001$, $t(20)=25.30$,<br>$d=5.52$ |
| Space Invaders | $p < .001$, $t(22)=21.02$,<br>$d=4.38$ | $p < .001$, $t(22)=18.70$,<br>$d=3.90$ | $p < .001$, $t(22)=29.04$,<br>$d=6.05$ |
| Enduro | $p < .001$, $t(22)=24.92$,<br>$d=5.20$ | $p < .001$, $t(22)=20.91$,<br>$d=4.36$ | $p < .001$, $t(22)=30.20$,<br>$d=6.30$ |

Results of the one-sample t-test for testing the subject’s correlation coefficient above chance level for all three models and games. ‘d’ represents Cohen’s d.

A comparison between the games revealed that all three models achieved their highest prediction accuracy in the game Enduro (*r_BaselineDQN_* = .30*, r_Ape_*_−_*_X_* = .35*, r_SEED_* = .48, with *r_DQN_* denoting the Pearson correlation coefficient of the time series predicted by the DQN), compared to Breakout (*r_BaselineDQN_* = .16*, r_Ape_*_−_*_X_* = .26*, r_SEED_* = .33) and Space Invaders (*r_BaselineDQN_* = .20*, r_Ape_*_−_*_X_* = .17*, r_SEED_* = .28) (see Fig 2). To assess the impact of advancements in DQNs on predictive power, we used a one-way analysis of variance (ANOVA) with repeated measures to conduct a comparison of prediction accuracy between the baseline DQN, Ape-X, and SEED. There was a statistically significant difference between the three DQNs in Breakout (Fisher’s z-transformed, F(2,40)=270.46, p*< .*001), Space Invaders (Fisher’s z-transformed, F(1.47,32.3)=95.29, p*< .*001, Greenhouse-Geisser-corrected), and Enduro (Fisher’s z-transformed, F(1.30,28.65)=158.41, p*< .*001, Greenhouse-Geisser-corrected). A post hoc paired t-test revealed a statistically significant higher correlation of SEED compared to the baseline DQN and Ape-X in Breakout, Space Invaders, and Enduro (Fisher’s z-transformed, p*< .*001, Bonferroni-corrected). These findings demonstrate that SEED, the most advanced of the three DQNs, provided the most effective feature-generating mapping for the GLM. Furthermore, we observed that the correlation of Ape-X significantly differed from the baseline DQN in the games Breakout and Enduro (Fisher’s z-transformed, p*< .*001, Bonferroni-corrected) but not in Space Invaders (Fisher’s z-transformed, *p* = 0.062, Bonferroni-corrected).

**Fig 2.**
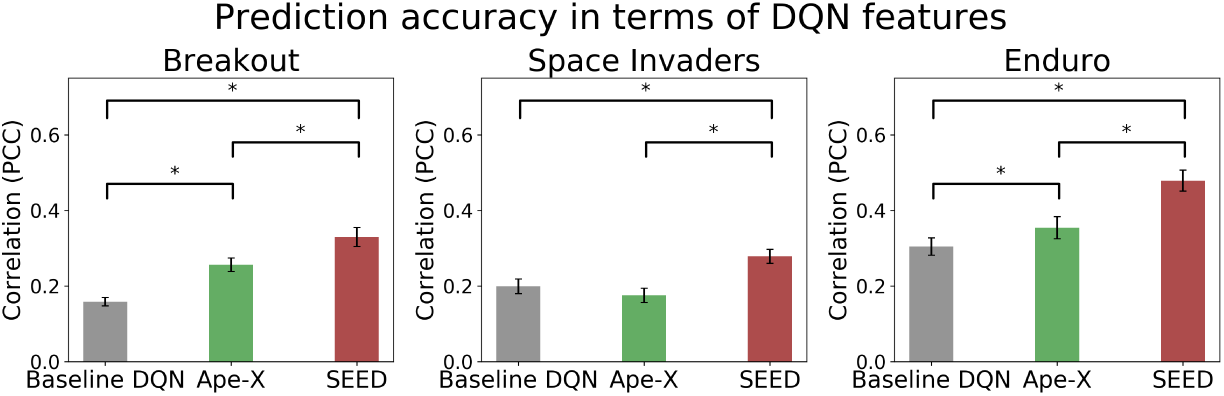
Prediction accuracies of the models. Pearson correlation of the three DQNs evaluated for Breakout (left), Space Invaders (middle), and Enduro (right). Error bars represent the 95% confidence interval for the correlation’s mean across subjects. The statistically significant differences between the DQNs (paired t-test, p*<*.001, Bonferroni-corrected) are denoted by ‘*’. Among the three models, SEED exhibited the highest ability to model human actions in all three games.

### 1.2 The balance between loss of information and predictive power: the optimal smoothing kernel

To analyze human gameplay behavior, the time series of human motor responses were modeled using the time series of DQN’s Q-values. However, examining individual frames of the time series did not seem meaningful for this kind of games due to the high temporal resolution of 45 Hz. Additionally, the DQN time series had a different structure, jitter, and characteristics due to the method of calculation, compared to those of humans. Hence, as an important preprocessing step, the DQN and the human time series were convolved with a Gaussian kernel 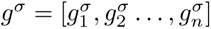 with a smoothing parameter *σ*, which is a measure for the strength of smoothing. Smoothing a time series *u* = [*u*_1_*, u*_2_*, . . . , u_m_*], the *k*-th element of the convolution vector *u* ∗ *g^σ^* is then defined by

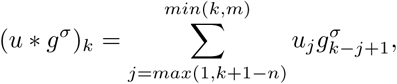

for *k* ∈ [1*, n* + *m* − 1]. This weighted averaging, where neighboring values are assigned greater weight based on their proximity, removes high-frequency components from the time series, making the time series smoother and more stable. In the standard cross-validation procedure of the GLM, we fitted the smoothed predictors to the smoothed human time series, both with the same smoothing parameter *σ* (see Fig 3 A). We found that the smoothing parameter significantly influenced prediction accuracy. In this section, we aimed to quantify the degree of information loss caused by smoothing, thereby determining the attainable temporal resolution when modeling human behavior. Therefore, we also analyzed the prediction of the unsmoothed, noisy human time series (see Fig 3 B). For this purpose, we calculated prediction accuracies as functions of the smoothing parameter *σ*.

**Fig 3.**
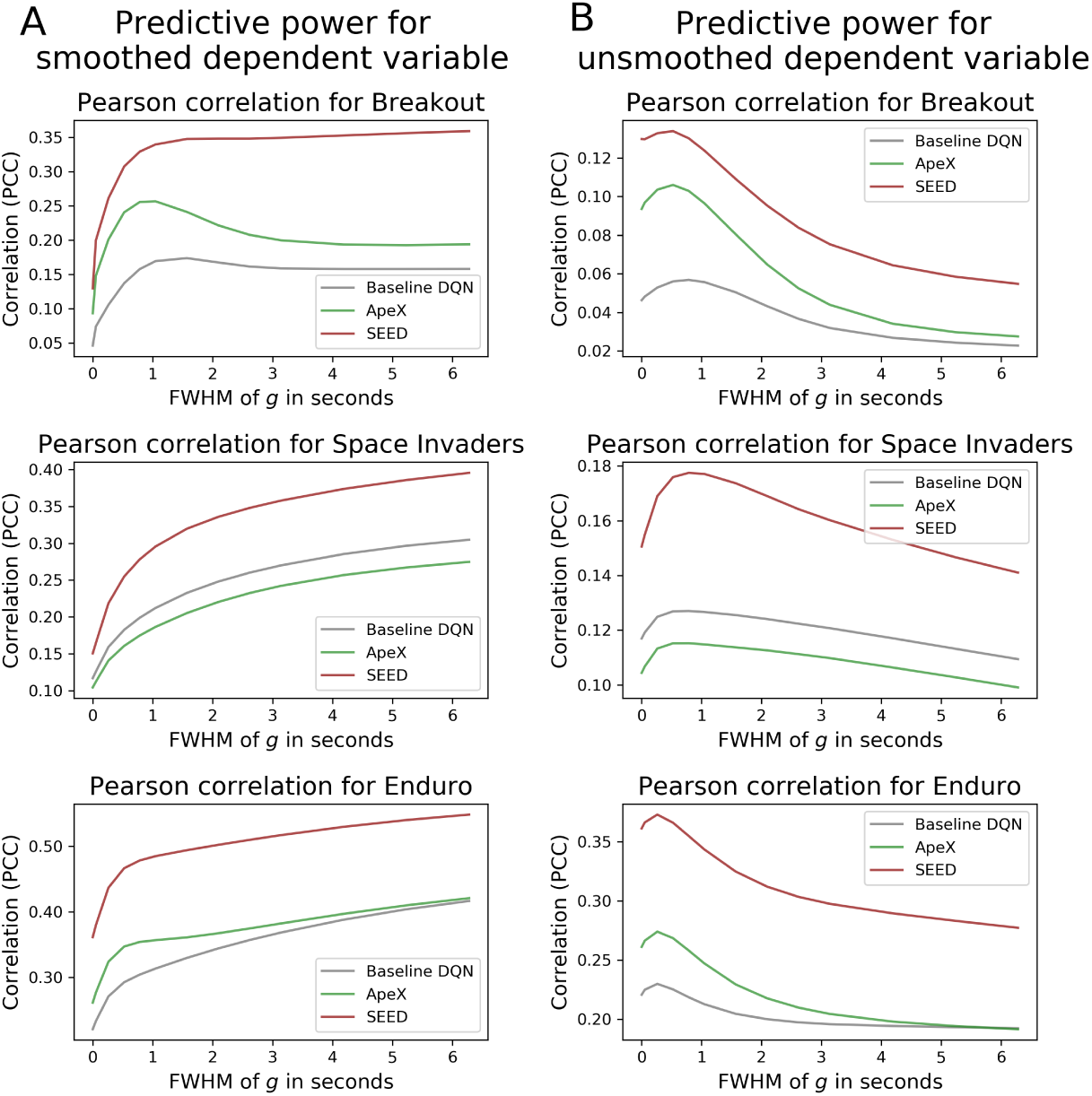
The influence of the smoothing parameter on prediction accuracy. The influence of the smoothing parameter on prediction accuracy. Graphs illustrate the Pearson correlation coefficient concerning the smoothing parameter of the Gaussian kernel *g^σ^*. A logistic model was fitted with its smoothed predictors in a cross-validation procedure to the smoothed human time series. Predictive power was assessed through Pearson correlation between the time series being modeled and the smoothed (A) and unsmoothed (B) human time series for Breakout (top), Space Invaders (middle), and Enduro (bottom) across the three DQNs. As the smoothing kernel increased, the prediction accuracy of the smoothed human time series increased (A). Conversely, as the information loss in higher frequencies in the predictors increased, the predictive power of the actual human time series decreased (B).

When predicting the smoothed human time series in the games Space Invaders and Enduro, it is noticeable that the more information removed in the high-frequency domain (more smoothing), the better the prediction. However, in Breakout the baseline DQN and Ape-X had already achieved the maximum predictive power when smoothing with a Gaussian kernel with FWHM = 1.57 seconds and FWHM= 1.05 seconds, respectively.

When predicting the original, unsmoothed human time series by smoothing the predictors using a Gaussian kernel with FWHM less than 0.79 seconds for Breakout and Enduro, and less than 0.26 seconds for Space Invaders, the prediction accuracy increased as the smoothing kernel increased. However, using a Gaussian kernel with a larger width led to a decrease in prediction accuracy.

An analysis of the data using fast Fourier transform (FFT) provided further insights into the effects of smoothing on prediction accuracy (see S7 Text). Smoothing the time series of the DQNs and those of humans resulted in a convergence of the frequency spectra of the predictors and the dependent variable. Such alignment may play a significant role in the observed increase in predictive performance. When predicting the original, unsmoothed human time series with its high frequencies, smoothing the predictors led to a divergence of the frequency spectra. This smoothing-induced loss of information in the high frequency domain of the predictors is possibly the explanation for why the unsmoothed human time series became challenging to model with progressive smoothing. Small Gaussian kernels initially had a positive impact on the predictive power of the actual human time series, as smoothing aligns the different structures within the time series, while also preventing overfitting during model fitting in the 7-fold cross-validation procedure. Overfitting could be attributed to the high frequencies present in the unsmoothed predictors, which captured the inherent noise in the unsmoothed human time series. This further highlights the importance of this preprocessing step.

To address this trade-off between information loss due to smoothing and reduced predictive power due to high temporal resolution, we opted for a Gaussian kernel with an FWHM = 0.79 seconds for data preprocessing of the human and DQN time series.

### 1.3 Prediction accuracy of DQNs relies on gaming performance

Before using DQNs as feature-generating mappings, they required several days of training to acquire the ability to play arcade games (see Fig 5). We hypothesized that training DQNs on progressively larger amounts of data, achieved through longer training times, would not only lead to higher game scores but also improve the predictive performance of the DQN. Throughout the training process, we stored multiple checkpoints for the baseline DQN and Ape-X, capturing the state of the DQNs at different time points. Since checkpoints were not available for SEED, our analysis exclusively focused on the baseline DQN and Ape-X. At each checkpoint, we evaluated the model’s prediction accuracy using the Pearson correlation coefficient between the actual human time series and the time series predicted by the model at that checkpoint (see Fig 4). We evaluated gaming performance using the average game score achieved by the DQN at that checkpoint.

**Fig 4.**
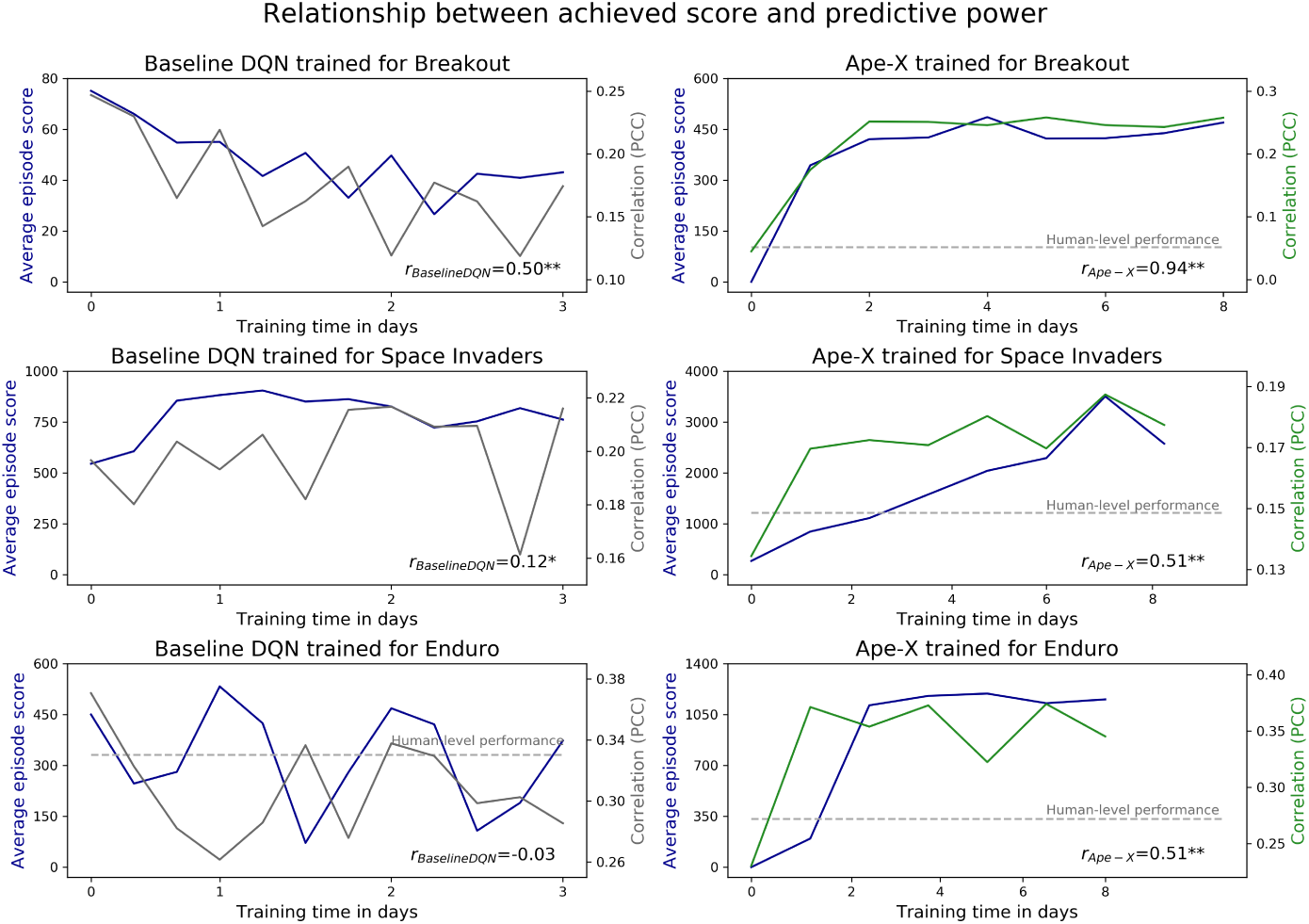
The influence of the training time on prediction accuracy. Relationship between the training time of a DQN, the gaming performance of a DQN, and the predictive performance of human time series using features generated by a DQN for Breakout (top), Space Invaders (middle), and Enduro (bottom). Each plot illustrates the learning curves (blue), showing the average game scores achieved in an episode, averaged across 50 episodes. Prediction accuracy is represented by the Pearson correlation coefficient for the baseline DQN (left, gray) and Ape-X (right, green). We calculated the Pearson correlation *r_DQN_* between gaming performance and predictive power as means across all subjects (lower right of each chart). Correlations significantly differing from zero were identified using one-sample t-tests, with ** p*<* 0.001 and * p*<* 0.01.

**Fig 5.**
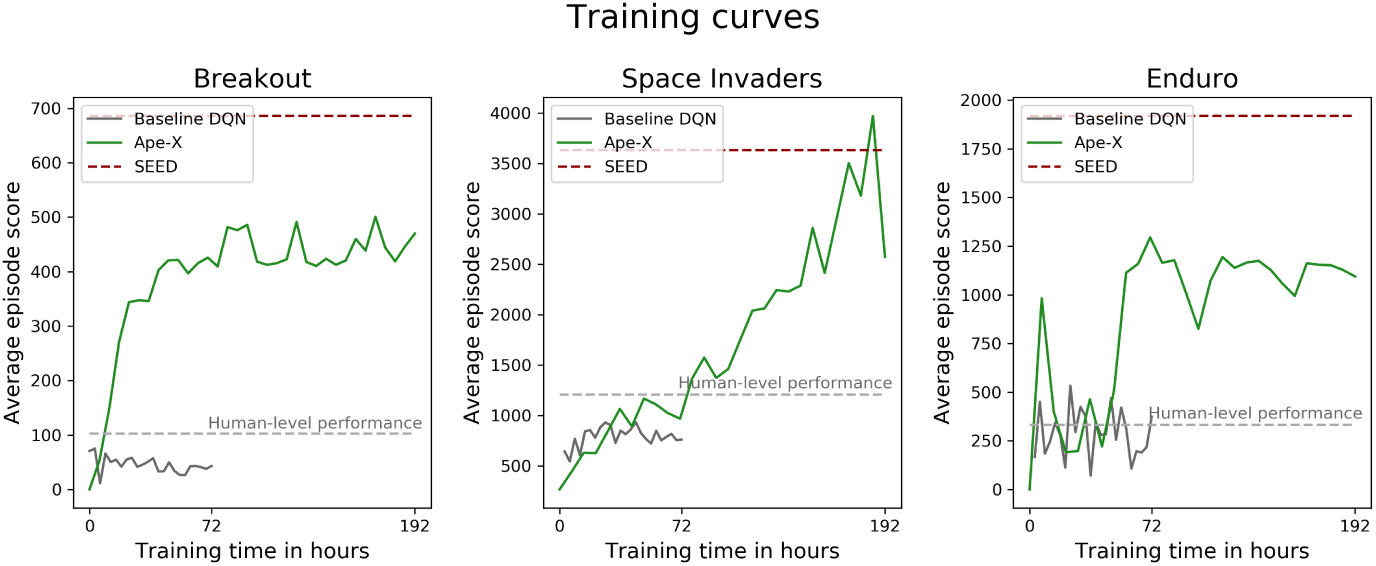
Learning curves of the DQNs. Learning curves for the baseline DQN (gray), Ape-X (green), and SEED (red). Each plot displays the game score achieved in an episode, averaged across 50 episodes per checkpoint across the training period and the human-level performance, represented by the average game score achieved by the participants per episode (light gray). Ape-X and SEED achieved human-level performance in all three games; the baseline DQN reached human-level in Enduro.

When examining the score curve trajectory of the baseline DQN, instability in the training process was observed. Therefore, an improvement in prediction accuracy with extended training time could not be expected and was not observed. A linear relationship between the average score achieved in an episode by the baseline DQN and its prediction accuracy was only evident in Breakout. In contrast, Ape-X showed significantly more stable learning curves that increased steadily, with occasional minor downward deviations as training time increased. However, the prediction accuracy is correlated with the gaming performance of Ape-X, confirming our initial intuitive hypothesis, which is particularly evident in Breakout. In Space Invaders, the model achieving the highest average score also reached its maximum predictive power. In Enduro, achieving a steady state in terms of performance does not necessarily correspond to a constant maintenance of prediction accuracy. However, it is somewhat surprising that even networks with low training duration and thus randomly initialized weights already exhibited positive correlations (see S2 Text).

## 2 Discussion

Modeling human behavior within complex and time-continuous environments has been challenging. However, recent advancements in machine learning, specifically the use of DQNs, have enabled the investigation of how humans process high-dimensional visual stimuli into appropriate motor responses in such environments.

In this work, we aimed to explore a DQN-based modeling approach for predicting human motor responses in complex and time-continuous tasks. The primary objective was to determine whether advancements in deep learning would translate into better predictions of human behavior. We evaluated the prediction accuracy of features generated by two advanced DQNs (Ape-X and SEED) in comparison to those of a baseline DQN when modeling human behavior during gameplay of three arcade games. Our results from the baseline DQN confirm existing evidence that a basic DQN model serves as a suitable tool for generating features for a GLM, enabling the modeling of game-based human behavior in arcade games [4, 5].

In recent years, DQNs have experienced a significant increase in power, reaching or surpassing human-level performance in various tasks [6, 11]. It seems plausible that advancements in machine learning can be leveraged to improve predictive capabilities [12, 13]. Our findings support this hypothesis. We demonstrated that SEED, the top-performing model, generated features that led to the most accurate predictions of human actions. We speculate that potential reasons for variations in prediction performance are mainly attributable to the differences in network architecture. Compared to the complex processes in the human brain, the baseline model and Ape-X being constructed with a feed-forward architecture and SEED featuring a combination of feed-forward and recurrent components, presumably represent significant simplifications [14]. We selected the DQN from [1] as our baseline due to its simple and fundamental architecture, as well as its human-comparable performance, making it a reasonable reference model. The learning curves of our baseline DQN implementation exhibited instability during training (see Fig 5), consistent with the results reported in the original paper [1] and confirmed by [15]. This alignment between our implementation and the original findings strongly suggests that the observed instability is an inherent characteristic of the algorithm and its architecture. Furthermore, this training behavior was a notable distinction from more recent models, demonstrating significantly more stable learning curves. The baseline DQN, a breakthrough in 2015, was the first to demonstrate that deep neural networks could learn to play Atari games directly from raw pixel inputs using reinforcement learning. This makes it all the more intriguing that, despite its simplicity, it achieved predictions above chance level. Yet, all three DQNs proved to be suitable for generating features to successfully model human behavior. A comparison of the three network architectures speaks to the importance and potential influence of certain network components, not only on game performance but also on predictive power. Ape-X and SEED additionally incorporated a dueling DQN architecture to distinguish between the value of actions and the value of the current state [16]. While this innovation might have contributed to the notable performance improvements in playing arcade games, it unexpectedly did not lead to a significant enhancement in the predictive power of the generated features of Ape-X in Space Invaders (see Section 1.1). Building upon the Ape-X architecture, SEED introduced a major advancement. In contrast to the feed-forward network, SEED, with its LSTM, employed an RNN, equipping the model with a temporal comprehension [17]. RNNs play a crucial role in modeling the human brain [14] and have been argued to contribute to the creation of a more human-like model [18, 19]. To evaluate the impact of temporal context on prediction accuracy, we additionally calculated the accuracy while disabling the pre-trained SEED’s LSTM from storing past game information (see S9 Text). The results show that without temporal context, SEED’s prediction performance was comparable to that of Ape-X and the baseline DQN. This suggests that the temporal context plays a crucial role in the model’s calculations and highlights the importance of this advancement for achieving high prediction accuracy.

Since DQNs first enabled the modeling of time-continuous behavior without manually designed features [4, 5], it would also be interesting to extend our hypothesis to a broader range of RL algorithms. The algorithms are categorized as model-based and model-free [22]. In our study, we used the model-free approach with Q-learning as the basis for comparing the three DQNs and their advancements. In addition to

Q-learning, policy optimization directly optimizes the agent’s policy. This algorithm has the advantage of applying to continuous action spaces, allowing for more complex tasks, whereas DQNs suit discrete actions [23]. While model-free algorithms require the optimal policy to be discovered through exploration, model-based algorithms have a functional representation of their environment and make decisions based on simulated outcomes. Model-free algorithms require significantly more training data than humans (see Section 3.2.3). Model-based algorithms can bridge this gap in Atari games, for example, by using stochastic video prediction techniques [24], though their performance remained relatively limited. We assume that similar learning efficiency enables insightful feature-generating mappings. A related approach is theory-based RL [25], which extends model-based approaches by incorporating human-like, abstract models of causal relationships. Since humans use internal models to plan and generalize in complex environments, the finding that the model-based algorithm EMPA can capture similarities with the brain’s model representation while learning Atari games [26] offers another interesting approach for feature-generating mappings. These RL algorithms can also be combined [27] as psychological and neuroscience studies showed their coexistence and interaction [28, 29]. Transformer models [21] and multi-task learning [20, 30] are expanding RL research, offering new ways to test our hypothesis.

The prediction accuracy relied not only on the DQN but also on the chosen smoothing kernel during data preprocessing. Smoothing with a Gaussian kernel was an essential preprocessing step in our analysis of time-continuous behavior. The Gaussian kernel acts as a low-pass filter, suppressing high-frequency components. This process reduces noise and makes trends more apparent. This step was crucial because conducting a frame-by-frame comparison at a resolution of 45 Hz did not seem meaningful. Despite the discrete implementation of the task, humans perceive tasks at such finely-grained resolutions as time-continuous [10]. However, at this temporal resolution, subjects were unable to perceive individual frames and thus could not execute a conscious motor response to each frame [31]. Furthermore, the DQN time series inherently differed in structure, jitter and characteristics from those of humans, as they exhibited high frequencies, for instance, due to the calculation of Q-values within the RL process (see S1 Fig), as well as the lack of a cost function, which led to unstable action selection and a jittery game flow (see S1 Video). As a result of smoothing, the time series became more stable for predictions, as the high-frequency components were removed. Therefore, in Section 1.2, we investigated the optimal smoothing kernel to ensure precise and meaningful predictions of human motor responses, while also maximizing the information content of the human time series. The impact of smoothing on the discrepancies and issues mentioned above can be observed in Fig 3. When the human and DQN time series were smoothed in parallel, the frequency spectra of both variables converged (see S7 Text), which could play a crucial role in the observed increase in prediction accuracy. For this reason, we were also able to show that smoothing the jittery DQN time series with an FWHM of 0.79 seconds was necessary to optimally predict the actual human time series, while also preventing overfitting during model fitting. Since predicting high frequencies is generally more challenging, the results demonstrate that smoothing a valid approach. The correlation peaked for a Gaussian smoothing kernel with FWHM = 0.79 seconds, demonstrating that human behavior can be modeled on a fine-grained temporal scale. This introduces a novel and important aspect, raising a key question in the analysis of time-continuous studies, which is relevant not only in the context of arcade game environments but also in time-continuous experimental tasks in general.

The use of DQNs represents a modern method for addressing questions in cognitive science [32–34]. However, there exists a scientific debate regarding whether DQNs can serve as models for cognitive processes due to the similarities and disparities they exhibit compared to the human brain [14, 35]. The degree to which a DQN can effectively serve as a model to explain human behavior and cognitive processes remains also controversial [2, 14, 36]. In the following, we will address these points, focusing on key aspects relevant to our research.

While the brain serves as a biological blueprint for artificial neural networks, and the networks attain human-level performance across a wide range of tasks, there are various disparities between the human brain and neural networks, such as differences in training behavior and architecture [14, 35, 37]. For example, humans possess the ability to interpret objects depicted on screens and leverage prior knowledge about the temporal and spatial properties of these objects [38, 39]. They could use their knowledge of cars in Enduro, for instance, to efficiently learn the effects of the actions ‘steering right’, ‘braking’, and ‘accelerating’. In contrast, our feed-forward DQNs do not possess these processes of cognitive abstractions inherent in the human brain and must acquire outcomes through stimulus-response learning.

However, DQNs also leverage certain advantages. During the training process, they process information from millions of frames [8] within a few days of training, a task that surpasses the temporal capacity of humans. To compensate for this in our main experiment, participants were given an informational advantage before the training period, including details about the game’s objective, the scoring system, and available actions and their effects in the game. The DQNs were also unaffected by factors like fatigue [40] or response time [41]. Therefore, it would be interesting to equip the DQN with constraints that mimic human limitations and investigate whether this has positive effects on predictive power. For example, restricting a DQN’s response time to human levels resulted in a decrease in game performance [42], but whether this translates to an increase in predictive power is still unclear. In our experiment, adjusting the human time series to match the response times of the DQN through a post-shift of the human time series did not lead to an improvement in prediction (see S6 Text).

All these features of DQNs led to superhuman performance as their complexity increased. Intuitively, one might assume that proximity to human-level performance could also be associated with increased predictive power. However, as human-level performance was significantly lower than that of Ape-X and SEED and comparable to the baseline DQN, the results in Fig 2 and Fig 5 speak against this intuitive assumption.

While Fig 4 demonstrates a positive correlation within Ape-X between a DQN’s task-solving ability and its prediction accuracy, it also highlights that upon reaching human-level performance, the prediction accuracy remained significantly below that of a model checkpoint with superhuman performance. Additionally, even highly efficient players, who might be expected to align closely with optimal strategies, did not show higher prediction accuracy when using modern DQNs (see S8 Text). This confirms that SEED’s high predictive power cannot simply be explained by its higher game scores. Achieving human-level scores was not sufficient to generate suitable features for prediction (see Fig 2, Fig 4, and Fig 5). This may be due to differences in the intrinsic processing and calculation of Q-values compared to human behavior. For this reason, we aimed to test this hypothesis empirically in our study. Our results demonstrate that despite the large gap in the game scores achieved by SEED and humans, SEED generated features that closely align with the representation of human behavior. While human factors such as fatigue, reaction time, and attention contributed to performance differences, this does not necessarily imply that DQNs extract different features from the input and weight them differently compared to subjects.

DQNs are primarily designed to automate complex tasks across diverse domains, focusing on rapid training on large datasets rather than explicitly modeling human brain processes [14, 35, 43]. Studies have shown that adapting the DQN to aspects such as human visual attention not only enhanced task performance in playing arcade games but also improved action prediction accuracy [44, 45]. In our experiment, we trained the DQNs without human data and did not instruct them to emulate human behavior. Although the DQNs we used received the same input as the participants, consisting of 84 × 84 grayscale screens, and the DQNs and the human participants shared the goal of maximizing game scores, their gaming behavior in reaching this goal differed. This became apparent when we visually analyzed the DQN behavior (see S1 Video) and when categorizing the gaming behavior in comparison to that of humans. At first glance, Ape-X and SEED exhibited gaming behavior patterns that resembled those of humans. However, upon closer visual investigation of the gameplay, disparities in the control mechanisms and the selection of actions became evident, deviating from human gameplay behavior. For example, the DQNs did not account for internal costs that subjects associated with executing an action (see S1 Text), resulting in arbitrary movements in “free choice” situations. In contrast, humans showed a tendency to perform no action in such situations [46]. One approach to address this divergence could involve the incorporation of regularization by penalizing the selection of an action other than ‘noop’ with a small negative reward. In Enduro, each action executed at any time step influenced the progression of the game, whereas, in Breakout, the paddle’s control mainly affected gameplay only when the ball was near the paddle. Given Enduro’s significantly higher predictive power across all three DQNs, we speculate that implementing the cost function in the DQN algorithms could enhance prediction accuracy.

When observing SEED’s gaming behavior, we noticed that it primarily followed specific strategies. For instance, in Breakout, it aimed to target the high-scoring top layer of the wall as early as possible, while in Space Invaders, it consistently aimed at the bonus spaceship. We observed similar strategic behavior in human gameplay. However, this behavior was not observed in the other two models, suggesting that reward clipping represents an additional technical limitation. Reward clipping is a feature implemented in the baseline DQN and Ape-X to stabilize the training process by restricting the received reward to a certain range, preventing the agent from choosing high-value targets [47]. In addition to the visual comparison of gameplay behaviors, analyzing the response patterns of the DQNs could offer deeper insights into the internal computations and game strategies that allow SEED to reach a feature space closer to modeling human behavior, improving prediction accuracy. To better understand which aspects of human action selection are captured by the DQNs, a valuable avenue for future research would be to develop an analytical approach for exploring similar state-space representations and examining the response behaviors within them.

While certain proposed adjustments to enhance the human-likeness of DQNs might enhance prediction accuracy, our study did not aim to implement a DQN that mimics human behavior. Rather, our goal was to demonstrate that DQNs can generate stimulus features that serve as suitable predictors in the GLM for modeling human behavior.

When dealing with neural networks, a second debate arises regarding the evaluation of DQNs as cognitive process models [2, 48] and their explanatory properties [2, 36]. Can black boxes, such as DQNs, effectively explain other black boxes, such as the human brain? The more complex DQNs become, the more opaque they tend to be. That the information obtained from neural networks can provide relevant insights into cognitive processes has become evident through a growing body of research that has addressed the comparison of representations of stimuli in neural networks with those in the human brain [14, 49–51]. For instance, the mechanisms underlying human brain-based planning and decision-making remained largely unclear. Therefore, neural network algorithms offer a potential avenue for modeling how the planning processes in the human brain might be instantiated [52]. Especially in our case, when comparing DQNs of varying complexity regarding their predictive capabilities, we aimed to gain insights into the respective strengths and limitations of each approach and the possibility of identifying an additional explanatory facet that could be ascribed to these models [2]. A systematic comparison of the architectures, such as evaluating the predictive power with and without individual components like the LSTM or dueling Q-network, and quantitatively assessing the importance of each exceeds our computational resources, as the DQNs would need to be retrained. For example, SEED was developed at Google with a large GPU cluster (e.g., a TPU pod with 2048 cores were used), which is also why we had to use a pre-trained SEED model. Due to our limited resources, replicating this is not feasible for us, even though it could provide exciting new insights. Therefore, we compared and interpreted our results in the context of findings from other studies. However, focusing solely on behavioral data in this study has its limitations. While behavioral data provide valuable insights into behavior, they do not address the underlying neural processes and mechanisms. Incorporating additional neuroscientific measurement techniques, such as EEG or fMRI, could enhance this approach by investigating the temporal and spatial visual brain representations [5, 33]. By integrating the hidden layers of a DQN, these methods could offer a fine-grained characterization of the individual steps of stimulus-response transformations, potentially providing deeper mechanistic insights. The prevailing direction is characterized by ongoing improvements of these models while simultaneously developing techniques to enhance their transparency [53, 54] and exploring avenues for making them even more human-like [35, 48]. All these approaches make DQNs suitable models for processes in the human brain. Hence, we see great potential in this modeling approach.

In addition to conventional experiments, characterized by their trial-based study designs and the use of categorial, low-dimensional stimulus spaces, our approach presented here can expand the possibilities for investigating human behavior in time-continuous and complex environments. This DQN-based modeling approach has also been applied to address other questions: Neural networks have also provided detailed insights into neural activations within task-related brain regions while subjects played arcade games, using functional MRI data [5, 26]. Motivated by our comprehensive validation of the DQN-based modeling approach within a time-continuous framework, an interesting question opens up whether the hidden layers of these advanced DQNs can be used to obtain more accurate predictions of the activations of brain regions involved in stimulus-response transformation during relatively complex behaviors, such as gameplay based on neuroimaging data. We aim to investigate this question in future work.

## 3 Methods

### 3.1 Data collection

#### 3.1.1 Ethics statement

The experimental protocol was approved by the Ethics Committee of the TUD Dresden University of Technology (EK 359072019) and conducted in accordance with the Declaration of Helsinki. All participants gave written consent prior to their participation in the study and received either financial compensation or course credit for their participation.

#### 3.1.2 Participants

The participants were healthy adults (N=23, 15 female, 8 male; mean age: 24.7 years, age range: 20-34 years). All participants were right-handed and had normal or corrected-to-normal vision. Most of them were recruited from the student population at TUD Dresden University of Technology.

#### 3.1.3 Structure of the experiment

Each participant attended three appointments on three separate days. On each day, one of three arcade games - Breakout, Space Invaders, or Enduro - was played. The order in which the games were played was randomized among subjects. Each main experiment consisted of 7 sessions, each session lasting 7 minutes of gameplay, corresponding to 20,000 frames. During each session, multiple game episodes could be played. An episode refers to a single playthrough of a game from start to game over. Before gameplay, subjects received detailed instructions about the game, explaining the game mechanics, finger placement, objectives, and scoring. Following this, they practiced the game for 5 sessions. Participants who reached a game-specific, predetermined threshold value proceeded with the main experiment, while those falling below the threshold completed additional training before moving on. Data from participants failing to meet the threshold, even after additional training, were excluded, resulting in *N* = 21 participants for Breakout and *N* = 23 participants for Space Invaders and Enduro.

Subjects were compensated with 10 EUR/h. In the main experiment, participants were rewarded with an additional linear gain proportional to their game score (for Breakout 0.4 ct per point, for Space Invaders 0.4 ct per 10 points, and for Enduro 0.4 ct per point). Between sessions, subjects were able to see the amount they had already won.

#### 3.1.4 Presentation of the task

The experiment was conducted on a computer running the Ubuntu 20.04 operating system. The experimental framework, including game instructions, was presented using Python’s TKinter library. To run the games and record participants’ data, we utilized the Arcade Learning Environment (ALE, [55]) and a modified Python script of the ALE Python interface [56], originally written by Ben Goodrich. We adapted and customized the framework to suit our study design. We used the original grayscale screen provided by the ALE library (ale.getScreenGrayscale), where a single game screen initially measured 160 × 210 pixels (width× height). Subsequently, we resized this screen to an array with 84 × 84 pixels to ensure consistency with the screen used by the DQNs (see Section 3.2.2). The game was displayed in full-screen mode on a 1280 × 1024 screen (width × height). To prevent flickering, we displayed the maximum value between the current and previous frames. To enhance readability for participants, we increased the resolution of the scoreboard. In Enduro, we reconstructed the scoreboard at a higher resolution. Additionally, we adjusted the contrast to improve the overall gameplay experience. Individual frames were presented at a frequency of 45 Hz.

#### 3.1.5 Arcade gameplay

The Atari 2600 games Breakout, Space Invaders, and Enduro were selected to cover a broad spectrum of gaming concepts. In Breakout, players are tasked with moving a paddle along the bottom of the screen to bounce a ball off it and break a wall of bricks at the top of the screen. Smashing a brick earns points, but missing the ball costs one of five lives. In Space Invaders, players control a fire cannon to prevent aliens from invading Earth, earning points for each alien destroyed while losing one of three lives if an alien hits the player’s cannon. In Enduro, players participate in a car race. The game aims to overtake 200 cars in the first round to progress to the second round, where the target increases to 300 cars. For each subsequent round, the player must overtake 300 cars to continue. Players earn a point for each car they successfully pass during the race. If the required number of points is reached before the day’s end, no more points can be earned for that day; failing to overtake enough cars ends the game episode. Response options include ‘no action’, ‘move left’, ‘move right’, ‘hit brake’ (only used in Enduro), and ‘fire’. Combinations of these actions are available in Space Invaders and Enduro. Participants used a German keyboard for input, using both hands. Control keys, ‘s’, ‘c’, ‘m’, and ‘l’, were marked with colored dots. In all games, the right middle and index fingers handled left (’m’) and right (’l’) movements. In Enduro, the left middle and index fingers were used to accelerate (’s’) and hit the brake (’c’), respectively. In Breakout, the left middle finger introduced a new ball into the game (’s’), and in Space Invaders, this finger was used to fire; the ‘c’ key had no function. To enhance paddle control in Breakout, we adjusted the steering (left and right). This involved executing a steering movement only on every second frame, with a ‘no action’ performed on the skipped frame, resulting in a deceleration of the paddle movement.

#### 3.1.6 Recorded data

Motor responses, received rewards, the number of episodes played in a session, and screen videos were recorded. The observed screens were stored as a sequence of matrices of pixel intensity values. The data were stored in a frame-based manner, with each frame corresponding to a data point, recorded at a rate of 45 Hz. Only the data from the main experiment were used for analysis.

### 3.2 Analysis of the data

#### 3.2.1 DQN-based modeling approach and prediction evaluation

We conducted a comparison of the prediction accuracy of features generated by three DQNs. Our analysis comprised two main components: the DQN itself and a GLM (see Fig 1). The DQN served as a nonlinear, feature-generating mapping, linking the visual stimuli to activations of neurons in its output layer (see Section 3.2.3). For each subject and each game, the human-generated videos were preprocessed (see Section 3.2.2), followed by processing the videos through a DQN specifically trained on that game, resulting in a time series of Q-values for each possible type of action. Before using the data in a GLM, the DQN-generated time series and the recorded human time series of motor responses were preprocessed (see Section 3.2.4). For each DQN, each game, each subject, and each type of action, we separately employed a logistic model to fit the generated stimulus features to human data. The time series of Q-values of a DQN served as predictors, while the human time series served as dependent variables. Within the design matrix, a time series of one neuron was replaced with a baseline of 1. Prediction accuracy was evaluated using a 7-fold cross-validation procedure. The DQN-generated time series of Q-values were fitted in the logistic model to human actions using six of the seven session blocks, with prediction evaluated on the left-out session block. With the calculated regression coefficients from training, the human time series of the left-out session was predicted using the generated Q-values as predictors. The Pearson correlation coefficient between the predicted and actual human time series of the left-out session was calculated. This process was repeated until each session block was used once as a test block.

To aggregate data, mean values were calculated across different variables. For each DQN and each game, averages of correlations across test blocks, subjects, and action types were computed. In Enduro, the action ‘break’ was excluded from consideration due to its infrequent use, only occurring in 50 out of 161 sessions (see S3 Text). Given the varying predictive power of different training checkpoints within a DQN (see Section 1.3), additional averages were determined in Sections 1.1 and 1.2 over the last four checkpoints of the trained models.

#### 3.2.2 DQN data preprocessing

To obtain the predictors for a GLM, the human-generated video screens were processed by a DQN. All three DQNs took a stack of four grayscale screens as input. Therefore, the grayscale video data were preprocessed. A maximum was computed over every third and fourth frame of the original ALE 45 Hz video, resulting in a video with a frequency of 11.25 Hz. The resulting frames (210 × 160) were then downsampled to a resolution of 84 × 84 using the Python commands skimage.transform for the baseline DQN and Ape-X, and cv2.resize for SEED. At each new time step, a current preprocessed frame was appended to the stack, while the oldest frame was removed from the stack (following a first-in, first-out approach).

#### 3.2.3 Tested DQN frameworks

We used three distinct DQN frameworks as feature-generating mappings for the GLM. Among these DQNs, the baseline DQN [8] represented the basic architecture and served as the baseline for comparison. The second DQN implemented was Ape-X [9], while the most advanced DQN tested in this study was SEED [7]. They all shared a common basic structure, consisting of three convolutional layers, with the rectified linear unit (ReLU) used as an activation function. The first hidden layer applied 32 filters with a kernel size of 8 and a stride of 4, the second layer applied 64 filters with a kernel size of 4 and a stride of 2, followed by the third layer with 64 filters, a kernel size of 3, and a stride of 1. These neural networks took a stack of four preprocessed frames (see Section 3.2.2) as input, with SEED additionally receiving information about the acquired reward. In the baseline DQN and Ape-X, each neuron in the output layer corresponded to an action of the game (as obtained from ale.getMinimalActionSet). In contrast, the output layer of SEED comprised 18 neurons, corresponding to all possible actions in the Atari environment, including those that did not affect the game (ale.getLegalActionSet). Consequently, this difference also affected the number of predictors in the respective GLM. The baseline DQN, Ape-X, and SEED varied in network architecture, training procedures, and model complexity while operating within the same gaming environment to predict human behavior. Below is a summary of each model:

##### Baseline DQN

We implemented a ‘vanilla’ feed-forward architecture consisting of three convolutional layers, followed by a fully-connected hidden layer consisting of 512 units, followed by the output layer. The algorithm used experience replay [57], a mechanism in which past gameplay experiences are stored in a buffer, allowing more efficient training by updating the network with a random subset of experiences at each step. The algorithm used a mini-batch size of 32. The discount factor *γ* was set to 0.99. The optimizer employed was torch.optim.Adam with a learning rate of 0.00025, and the loss function used was torch.nn.SmoothL1Loss. Additionally, an *c*-greedy algorithm was implemented with *c* = 0.05, balancing exploration and exploitation by allowing occasional random actions while prioritizing the learned policy. This model did not incorporate any additional features. The DQN received input consisting of the frames observed by humans and information about the game’s termination. Our code is based on the theory from [8].

##### Ape-X

This section is a summary of the key points from [9]. The algorithm showed that generating additional data through a distributed architecture and selecting it in a prioritized way was sufficient to improve the performance of deep RL models significantly. The Ape-X DQN was built upon the baseline DQN with several modifications aimed at improving performance. A dueling network architecture [16] refined the estimation of the Q-function by separately learning the estimation of the state value, and the estimation of action advantages, dependent on a state. Therefore, after the three convolutional layers, the network split into two streams for the two estimations: to estimate the state value, a fully connected layer with 512 units was used, followed by another fully connected layer that produces a single scalar value. To estimate the action advantages, a fully connected layer with 512 units was followed by another fully connected layer with a number of units corresponding to the possible actions in the game. These two streams are then combined in an output layer to calculate the Q-values. The DQN received input consisting of video screens and information about the game’s termination. The training process of the DQN was divided into two main parts: acting and learning. Here, an extended prioritized experience replay [58] was applied in the distributed setting. Multiple actors (four in our case, originally 360) with different policies collected a diverse set of data by exploring the environment, evaluating a policy, and depositing their experiences into a shared experience replay memory. Initial priorities were also calculated for these data to identify the most useful experiences. Subsequently, the learner updated the network by sampling batches of these prioritized data from the buffer and updating the priorities in the shared memory. These two processes ran concurrently. The learner’s network parameters were periodically used to update the actors’ networks. The learning algorithm was optimized using double Q-learning [59], which mitigated the overestimation of action values by decoupling action selection from action evaluation, combined with multi-step bootstrap targets [60]. Our implementation of Ape-X was based on the GitHub repository [61]. Modifications were made to adapt it to our specific data analysis pipeline, including adjustments related to data preprocessing and integration with the Atari environment. We used the following hyperparameters in the implementation: The discount factor was set to *γ* = 0.999. The *c*-greedy algorithm was employed with different values of *c* for the four actors: *c* = 0.04, *c* = 0.047, *c* = 0.0056, and *c* = 0.00066. The optimizer used was torch.optim.RMSprop with a learning rate of 0.00025*/*4. A mini-batch size of 50 was applied. The actors’ networks were updated every 200 iterations with the latest network parameters from the learner.

##### SEED

This section is a summary of the key points from [7]. SEED is an enhanced DQN that incorporates techniques from policy gradient methods [62] in addition to the Q-learning algorithm. SEED benefited from several advancements compared to Ape-X. One of the key improvements was implementing of a long short-term memory (LSTM) [17] in its architecture. LSTMs, being RNNs, can store information over longer periods, enabling them to consider contextual information in future decision-making processes. The LSTM regulates what information is stored, retained, or forgotten through various ‘gates’. After three convolutional layers and a fully connected layer, the LSTM was applied. The output of the LSTM was then fed into the dueling DQNs for further processing and finally merged in the output layer. As in Ape-X, the training also consisted of two parts: the actor and the learner, but with a reallocation of tasks. The central unit was the learner, which ran three types of threads: inference, data prefetching, and training. In the inference step, the learner sampled actions from a batch of observations and other game information. The trajectories were later sampled by the data prefetching threads. The training thread computed gradients using TPU cores and updated all model parameters synchronously. Only one copy of the model was maintained, ensuring that any optimizations to the model immediately impacted the action selection. The actor’s task was to interact with the environment by executing the actions provided by the learner and sending the observations back to it. To use resources efficiently, an actor could run multiple instances of the environment simultaneously. This reallocation enabled several improvements, including a reduction in bandwidth between the actor and learner, an increase in speed, and a decrease in computational overhead by performing inference on TPUs. An optimized distribution system, flexible adaptation to workloads, and efficient resource utilization made it possible to train on millions of frames per second. To minimize latency in communication between the learner and actor, a remote procedure call (gRPC) framework was introduced. Additional algorithms were implemented to stabilize and optimize the training process, including double Q-learning [59], multi-step bootstrap targets [60], prioritized replay [58], value-function rescaling [63], and V-trace [62]. The following parameters were used for SEED: 610 actors, *γ* = 0.997, a mini-batch size of 64, no reward clipping, an Adam optimizer from TensorFlow with a learning rate of 10^−4^, and *c*-greedy exploration with the *i*-th actor ∈ [0*, N* ) using 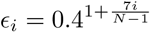. Additional parameters can be found in [7].

The implementation of SEED was based on the code repository available at [11]. Due to hardware limitations, training SEED from scratch was not feasible, thus pre-trained checkpoints provided by Google DeepMind were used. In addition to the observed screens, which served as input to the DQN for calculating the Q-values, supplementary input vectors were provided to the LSTM, including the reward vector and a vector denoting the game’s termination. In the original code, the LSTM received a vector representing the actions taken in the previous time step. To prevent potential bias from the action vector’s influence on feature generation and predictive power validation, it was fixed as a zero vector (indicating ‘no action’) during feature generation. However, this modification did not noticeably impact predictive power (see S5 Text), and SEED still achieved superhuman levels in playing arcade games.

The DQNs were trained via reinforcement learning separately on each of the games, without using any human data (see Fig 5). Training for the baseline DQN was terminated after three days, corresponding to approximately 60 million environmental frames. Ape-X underwent training for eight days, corresponding to approximately 800 million frames. Throughout the training process, the neural network weights were periodically locked into a fixed state and saved as checkpoints. During gameplay, the DQNs selected actions based on the highest estimated Q-value in a given state, using the *c*-Greedy algorithm.

#### 3.2.4 GLM data preprocessing

Before using the time series of Q-values as predictors and human actions as the dependent variables in the GLM, they had to be preprocessed. Due to the different sampling frequencies of human data (45 Hz) and DQN data (11.25 Hz), the Q-values required temporal upsampling to a common space, with each Q-value replicated four times within each time step. This resulted in DQN time series with a length of 20,000 data points for every processed video. For each time step, the Q-values were standardized across the actions by converting the networks’ top layer activations into z-scores. The human time series of motor responses were coded binary, indicating whether the action was performed at a given time step (coded as 1) or not (coded as 0). All combined actions, such as ‘fire-right’, ‘accelerate-left’, and ‘break-left’, were split into their action vectors: ‘fire’ and ‘right’, ‘accelerate’ and ‘left’, and ‘break’ and ‘left’, respectively. To make the human time series and the DQN time series match after temporal downsampling, a time shift was necessary. To achieve this, we excluded the first 12 time steps and the last three time steps from the preprocessed Q-value time series and removed the first 16 time steps of the human response variables.

Since a comparison between individual frames did not seem meaningful due to the high temporal resolution of 45 Hz, the predictors and the binary human time series were smoothed using a Gaussian kernel. The preprocessed vectors representing human actions could then be interpreted as response probabilities. In Sections 1.1 and 1.3, the time series were convolved using a Gaussian kernel with FWHM= 0.79 seconds. In Section 1.2, we conducted a more detailed investigation into the optimal smoothing parameter. For the plot in Fig 3 B, we fitted the smoothed DQN to smoothed human training datasets. On the test dataset, we calculated the correlation between the original, unsmoothed human time series and the predicted time series.

#### 3.2.5 Software and Hardware

The baseline DQN, Ape-X, and SEED were trained and tested on arcade games using the ALE framework. ALE served as an interface for research in the field of RL applied to Atari 2600 games. We used the Python interface by Ben Goodrich (https://github.com/bbitmaster/ale python interface). For the baseline DQN and Ape-X, we used Python 3.8.8 with PyTorch on Debian “Buster”, for SEED Python 3.6.9 and TensorFlow on Ubuntu 18.04.5 LTS.

Our computational setup consisted of a NVIDIA RTX A4000 GPU with 16GB of memory, and an Intel Xeon W-2245 CPU with 8 cores and 64 GB of RAM.

#### 3.2.6 Data availability statement

All behavioral human data, along with the Q-values generated by the DQNs and the resulting files, are publicly available at https://osf.io/xqmjc/ (DOI: 10.17605/OSF.IO/XQMJC). The experimental task, DQN code, and analysis code used for generating the results can be found in the public GitHub repository at https://github.com/SHaberland15/Arcade DQN Research. We used Zenodo to assign a DOI to the repository: 10.5281/zenodo.14974341.

## Financial Disclosure

The authors acknowledge support by the Deutsche Forschungsgemeinschaft (DFG) grant 445383113. The funders had no role in study design, data collection and analysis, decision to publish, or preparation of the manuscript.

## Notes

https://osf.io/xqmjc/

https://github.com/SHaberland15/Arcade_DQN_Research

### Competing Interest Statement

The authors have declared no competing interest.

