## Supplementary material for "Advances in deep reinforcement learning enable better predictions of human behavior in time-continuous tasks": S1 Fig

**S1 Fig. Visualization of human time series and generated features** Visualization of the original and preprocessed time series of a randomly selected subject for the actions 'fire' and 'right', as well as the original and preprocessed time series of features generated by Ape-X representing the neurons corresponding to the actions 'fire' and 'right' during the first session of Space Invaders.

(A) Time series of features (Q-values) generated by Ape-X.

(B) Preprocessed time series of features generated by Ape-X, smoothed with a Gaussian kernel with FWHM = 0.79 seconds.

(C) Binary time series of the subjects' actions.

(D) Preprocessed time series of the subjects' actions, smoothed with a Gaussian kernel with FWHM = 0.79 seconds.

(E) Time series of human motor responses predicted by the GLM with features generated by Ape-X.

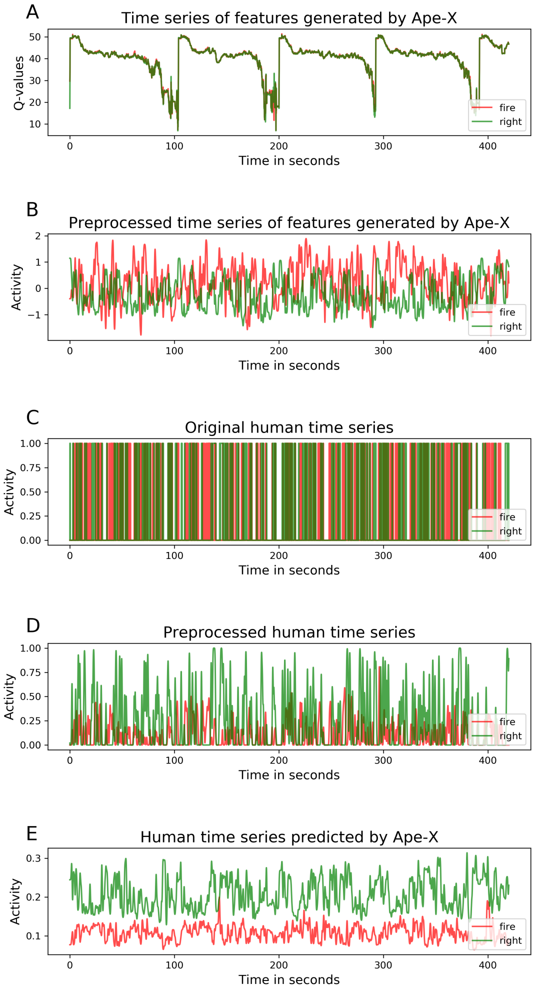
