## Supplementary material for "Advances in deep reinforcement learning enable better predictions of human behavior in time-continuous tasks": S1 Text

**S1 Text. Frequency of using the action 'no operation' by the DQNs and humans.** Humans compute internal costs associated with pressing a button. However, DQNs don't know this cost function. As a result, subjects tended to choose the 'no operation' action more frequently than DQNs. Participants responded to each frame of the game with an action, including the action 'no operation'. The videos were processed by a DQN, generating Q-values for each input screen and each possible action. DQNs selected the action with the highest Q-value. We compared the percentage of 'no operation' actions, relative to the total number of frames, between the subjects and the DQNs across all sessions and subjects for all three games. Subjects demonstrated a significantly greater tendency to refrain from pressing a button compared to the DQNs. The Baseline DQN has been shortened to B. DQN due to space constraints.

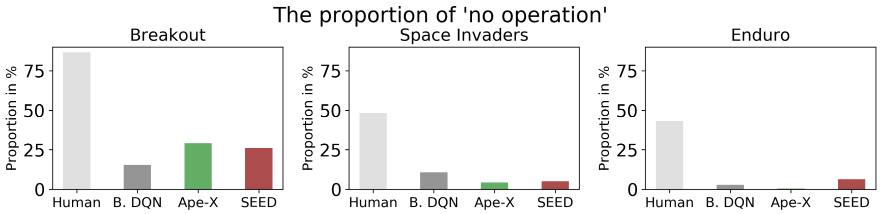
