## Supplementary material for "Advances in deep reinforcement learning enable better predictions of human behavior in time-continuous tasks": S2 Text

**S2 Text. The impact of DQN architecture on prediction performance.** The baseline DQN, Ape-X, and SEED shared a common fundamental structure: a feed-forward neural network consisting of three convolutional layers. The results of the training revealed that the features of the three DQNs possessed predictive power even after no or only a short training period. These results opened up the question of the extent to which architecture influences prediction. To investigate the potential underlying factors contributing to the initial positive correlations observed in the untrained DQNs, we evaluated the predictive power of features generated by two plain networks. The first network (linear NN) consisted of only two linear layers with the rectified linear unit (ReLU) as an activation function. The second network (conv NN) comprised a single convolutional layer with ReLU as an activation function, followed by a fully connected layer as an output layer. Importantly, these networks had no additional add-ons or the capability to play any of the Atari games. The weights of these networks were randomly initialized. The results were somewhat unexpected. These networks provided features that led to positive Pearson correlations in predicting human data, even without having learned specific information or patterns. This demonstrates that the fundamental network architecture of these models inherently possesses the ability to capture and generate features for predicting human responses. A similar observation has been made in the context of deep neural networks used for object recognition, where these networks were employed to study temporal and spatial visual brain representations [1].

**Pearson correlations for an untrained baseline DQN, an untrained Ape-X, an untrained SEED, an untrained fully connected neural network with two fully connected layers (linear NN), and an untrained convolutional neural network with one convolutional layer and one linear layer (conv NN) in Breakout, Space Invaders, and Enduro. All networks were initialized with random weights. The reported values represent the mean value of the Pearson correlation coefficient across 10 realizations per network.**

| | **Baseline DQN** | **Ape-X** | **SEED** | **linear NN** | **conv NN** |
|---|---|---|---|---|---|
| Breakout | 0.04 | 0.04 | 0.16 | 0.05 | 0.04 |
| Space Invaders | 0.14 | 0.16 | 0.28 | 0.14 | 0.15 |
| Enduro | 0.22 | 0.21 | 0.33 | 0.21 | 0.20 |
