## Supplementary material for "Advances in deep reinforcement learning enable better predictions of human behavior in time-continuous tasks": S3 Text

**S3 Text. Participants avoided braking in Enduro.** To evaluate prediction accuracy, for each game and each DQN we calculated the Pearson correlations across all subjects, sessions, the last four checkpoints, and action types. However, in Enduro, the action 'brake' was not used by participants in 111 out of 161 sessions, making predictions of this action impractical. Consequently, we excluded this type of action from the calculations.

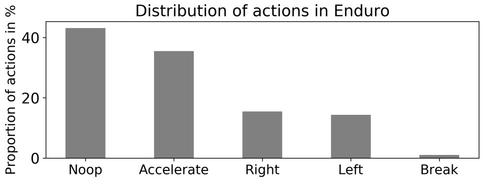


Each frame seen by the subjects required an action, including the action 'no operation' ('noop'). The bars depict the percentage of executions for each action type relative to the total number of frames in Enduro across all sessions and subjects. The plot revealed a notable disparity in the usage of the action "brake" compared to the other actions.
