## Supplementary material for "Advances in deep reinforcement learning enable better predictions of human behavior in time-continuous tasks": S4 Text

**S4 Text. Prediction accuracy for further investigations with fMRI data.** To further generalize the analysis of this approach to future studies using fMRI data, we re-evaluated the prediction accuracy using a Gaussian smoothing kernel with FWHM= 5.3 seconds. This value corresponds with the FWHM of the standard hemodynamic response function (HRF). Results comparable to those presented in the initial section of the Results are illustrated in this figure, employing the larger smoothing kernel.

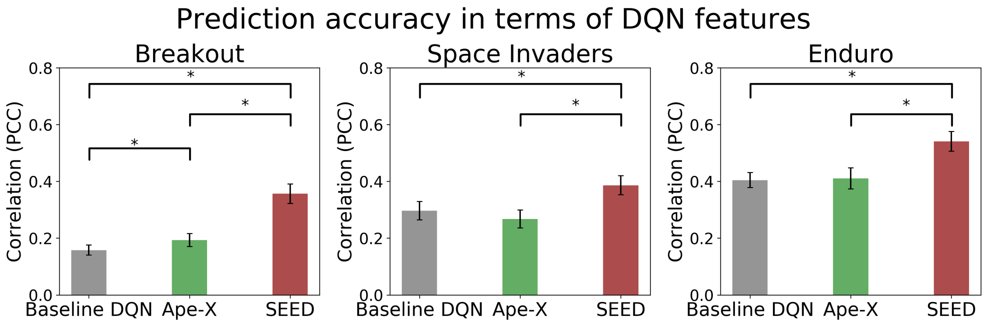


Pearson correlation of the three DQNs evaluated for Breakout (left), Space Invaders (middle), and Enduro (right). Error bars represent the 95% confidence interval for the correlation's mean across the subjects; The statistical significance ($p<.01$, Bonferroni-corrected, paired two-sample t-test) is denoted by '*'. Time series were smoothed using a Gaussian kernel with FWHM= 5.3 seconds.
