## Supplementary material for "Advances in deep reinforcement learning enable better predictions of human behavior in time-continuous tasks": S5 Text

**S5 Text. Prediction accuracy of SEED with and without code modifications.** In the original SEED code, an input vector representing executed actions was fed into the LSTM. However, to address the potential bias introduced by the action vector's influence on feature generation and the validation of predictive capability, we nullified this vector by setting its values to zero, corresponding to the action 'no operation'. A comparison of the predictive performance of features generated with the original code of SEED indicated that this modification had only a minor impact on prediction accuracy. However, significant effects between the DQNs remained.

**We manipulated the input of SEED's LSTM and compared the mean value of the Pearson correlation coefficient across the sessions, subjects, and types of actions for Breakout, Space Invaders, and Enduro with and without this modification. Time series were smoothed with a Gaussian kernel with FWHM=** $0.79$ **seconds.**

| | **Breakout** | **Space Invaders** | **Enduro** |
|---|---|---|---|
| Original SEED code | 0.29 | 0.30 | 0.46 |
| Modified SEED code | 0.32 | 0.27 | 0.48 |
