## Supplementary material for "Advances in deep reinforcement learning enable better predictions of human behavior in time-continuous tasks": S6 Text

**S6 Text. Incorporating human reaction times into prediction.** The response time of humans varies across individuals and is influenced by numerous factors, but averages around 250 ms for visual stimuli [1]. This corresponds to 11 frames at the temporal resolution of our experiment, which is 45 Hz, whereas a DQN typically has response times of only 1-4 frames. Therefore, we investigated a post-adjustment of the time series of humans and DQNs by shifting the human time series by 7 frames. However, this did not lead to a significant improvement in predictive power, contrary to expectations.

**Prediction accuracy, measured as the Pearson correlation coefficient, with subsequent consideration of human response times. We shifted the human time series by 7 frames to align with the time series generated by the DQNs. The time series were smoothed using a Gaussian kernel with FWHM**= 0.79 **seconds.**

| | **Breakout** | **Space Invaders** | **Enduro** |
|---|---|---|---|
| Baseline DQN | 0.16 | 0.20 | 0.31 |
| Ape-X | 0.25 | 0.18 | 0.37 |
| SEED | 0.32 | 0.28 | 0.48 |
