## Supplementary material for "Advances in deep reinforcement learning enable better predictions of human behavior in time-continuous tasks": S7 Text

**S7 Text. The impact of the smoothing kernel.** We investigated the trade-off between predictive power due to the high temporal resolution and information loss due to smoothing. We determined that the optimal smoothing kernel for modeling human behavior has a FWHM of 0.79 seconds. From the charts in Fig 3, it is observed that the effect of smoothing was influenced not only by the size of the smoothing kernel or the DQNs but also varied across different games, reflecting the inherent characteristics of the games within the dataset. To gain a deeper understanding of how smoothing affects the data, we performed a detailed analysis of the time series and their frequencies using the Fast Fourier Transform (FFT). In Fig 3 A, it is observed that prediction accuracy increased with a larger smoothing kernel. Since smoothing reduces higher frequencies within the time series, it caused the convergence of the frequency spectra among the predictors, the DQN time series, and the dependent variable, the human time series (see Fig A-D). Such alignment could contribute to the observed increase in predictive power. Simultaneously, this may explain why the actual unsmoothed human time series, with its high frequencies, became challenging to model as the predictors were progressively smoothed, retaining only low frequencies after sufficient smoothing (see Fig 3 B). The initial increase in the prediction of the actual, unsmoothed human time series in Fig 3 B by using smoothing kernels with a smaller width could be attributed to preventing overfitting. Eliminating high frequencies in the predictors might reduce the ability to fit noise present in the original human time series.

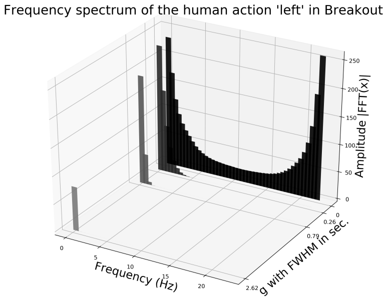


**Fig A.** Frequency-domain representation obtained through FFT applied to the time series representing subjects performing the action 'left' in Breakout. The FFT results were averaged across all subjects. Before FFT analysis, the time series were smoothed using smoothing kernels with FWHM of $[0, 0.26, 0.79, 2.61]$ seconds (different shades of green).

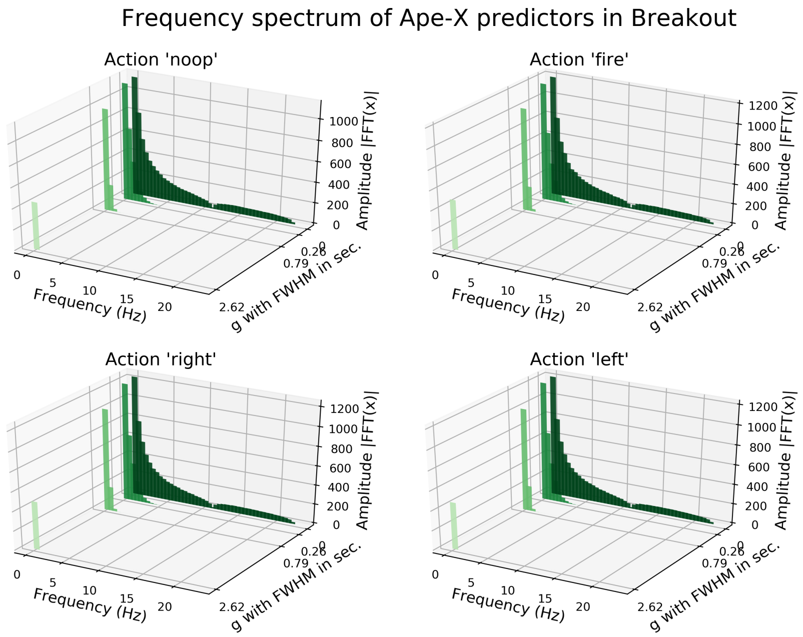


**Fig B.** Frequency-domain representations obtained through FFT applied to the predictors generated by Ape-X in Breakout. The FFT results were averaged across all subjects. Before FFT analysis, the time series were smoothed using smoothing kernels with FWHM of $[0, 0.26, 0.79, 2.61]$ seconds (different shades of green).

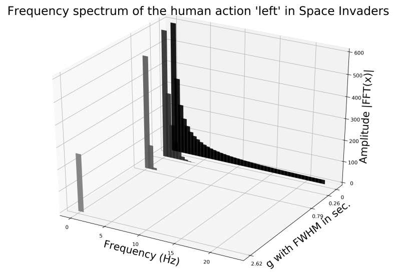


**Fig C.** Frequency-domain representation obtained through FFT applied to the time series representing subjects performing the action 'left' in Space Invaders. The FFT results were averaged across all subjects. Before FFT analysis, the time series were smoothed using smoothing kernels with FWHM of $[0, 0.26, 0.79, 2.61]$ seconds (different shades of green).

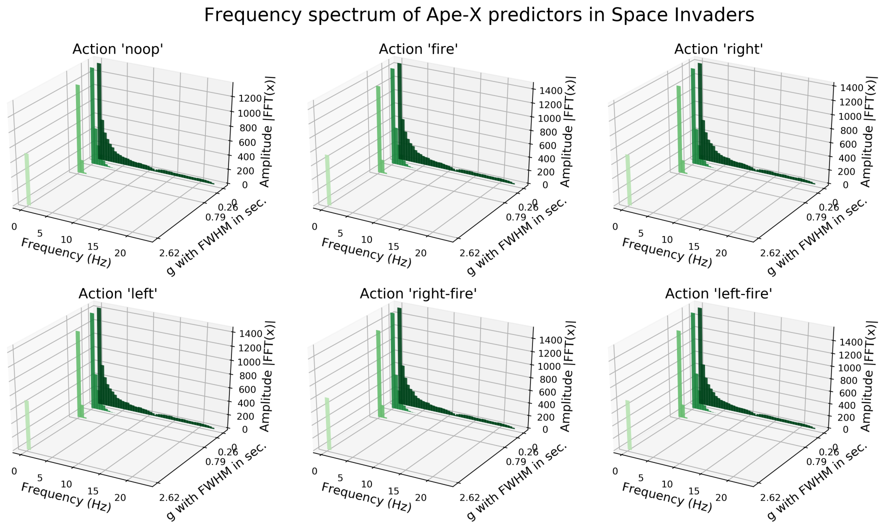


**Fig D.** Frequency-domain representations obtained through FFT applied to the predictors generated by Ape-X in Breakout. The FFT results were averaged across all subjects. Before FFT analysis,45 the time series were smoothed using smoothing kernels with FWHM of $[0, 0.26, 0.79, 2.61]$ seconds (different shades of green).

The effect of smoothing on prediction accuracy differed among games. We speculate that this variance could be due to differing compositions of frequency components within the human time series, which are probably influenced by the design of each game. For instance, the FFT analysis highlighted that the fast-paced game design of Breakout, characterized by actions executed in short intervals due to rapid steering and the modifications made to the controls (see Methods Section - Atari gameplay), was reflected in the presence of high frequencies in the human data (see Fig A).
