## Supplementary material for "Advances in deep reinforcement learning enable better predictions of human behavior in time-continuous tasks": S8 Text

**S8 Text. Relationship between subjects' gaming performance and DQNs' prediction accuracy** The DQN was trained to estimate the Q-value function, approximating the optimal policy for solving a task. As advancements in machine learning continue, DQNs get closer to this optimal policy. When playing arcade games with the goal of maximizing the game score, SEED achieved significantly higher scores than the baseline DQN and Ape-X. One might intuitively expect that subjects who perform better in arcade games are closer to the optimal gameplay strategy, and would exhibit greater prediction accuracy when using modern DQNs. However, as shown in the figure, this was not the case. One reason could be that even the best player among our measured subjects did not reach the level of SEED and Ape-X. Additionally, achieving human-level scores may not have been sufficient to generate suitable features for prediction (see also Fig 4). SEED generates features that closely align with representations of human behavior and demonstrates that similar scores do not necessarily indicate similar gameplay behavior.

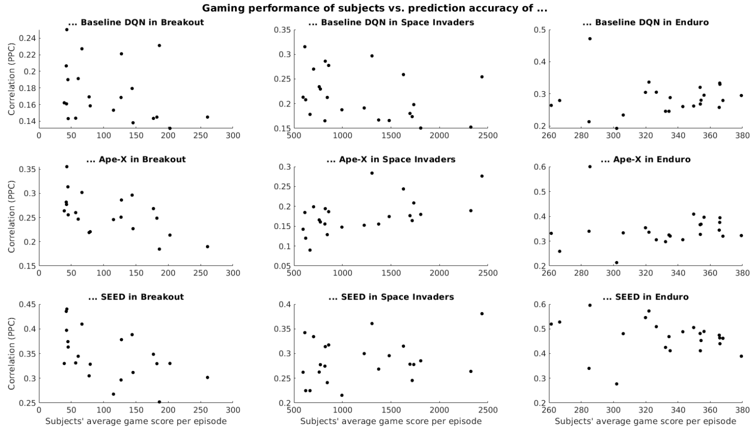


Scatter plots show the relationship between subjects' gaming performance and DQN prediction accuracy. Each point represents a subject, with gaming performance (average score across sessions) on the x-axis and prediction accuracy of the subject's time series on the y-axis.
