## Supplementary material for "Advances in deep reinforcement learning enable better predictions of human behavior in time-continuous tasks": S9 Text

**S9 Text. The importance of incorporating past experiences into current decision-making.** SEED, with its LSTM, can store information about the game state over longer periods, allowing it to retrieve contextual information with each new time step. The impact of this temporal aspect on decision-making is illustrated in the plot. We disabled the internal state of the recurrent cell in a pre-trained SEED by resetting the internal state at each time step. This means the model could not store information from previous screens, rewards, or actions, and each action calculation was based solely on the current input screen. The results show that our model with features generated by SEED and the original LSTM significantly outperformed the one with features generated by SEED and the modified LSTM (t-test, $p < 0.001$). Without the temporal context, however, SEED's performance was between that of the baseline DQN and Ape-X. This suggests that the temporal context played a crucial role in the model's calculations. However, since we could not retrain SEED with the modified LSTM input due to limited computational resources, the full impact of this factor for prediction accuracy remains an open question.

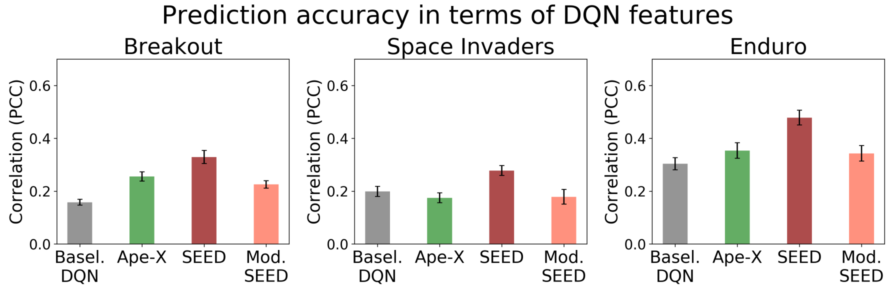


Prediction accuracy of SEED with deactivated LSTM state. Pearson correlation of the baseline DQN (gray), Ape-X (green), SEED (dark red) and SEED with modified LSTM (light red). Error bars represent the 95% confidence interval.
